# Receptor-guided AAV Tropism Engineering via MATCH

**DOI:** 10.64898/2026.03.31.715691

**Authors:** Nolan Graham, Satheesh Kumar, Joseph Rainaldi, Seoin Yang, Andrew Portell, Billy Santoso, Prashant Mali

## Abstract

Precise control over viral tropism remains a major challenge in the development of gene delivery technologies. We present MATCH (Modulation of AAV Tropism through Conjugation to Homing proteins), a modular biochemical method that enables programmable, receptor-guided retargeting of adeno-associated viruses (AAVs) through site-specific covalent protein conjugation. By incorporating a SpyTag peptide motif into selected AAV capsid loops, MATCH allows one-step, stoichiometrically defined attachment of recombinant SpyCatcher-linked targeting proteins to the viral surface. Using mosaic AAV-DJ and AAV9 capsids with controlled SpyTag incorporation, we achieve efficient assembly and tunable ligand display. MATCH-AAVs conjugated to an anti-CD3 single-chain antibody efficiently activate and transduce resting human T cells within mixed PBMC populations in vitro, achieving transduction levels of up to ∼58% of total PBMCs. Conjugation to transferrin receptor (TfR1)-binding proteins yielded enhanced brain transduction in vivo, with murine TfR1-targeted MATCH-AAV9 exhibiting up to an 84-fold increase in brain expression relative to wild-type AAV9. Human TfR1-targeted vectors similarly enabled robust, receptor-dependent transduction both in vitro and in humanized mouse models. Both TfR-targeted vectors enabled widespread transduction of the parenchyma, consistent with TfR1-mediated crossing of the blood-brain barrier. Finally, we establish a streamlined one-pot “Mix-and-MATCH” production strategy in which capsid and targeting ligands are co-expressed during vector generation, yielding functional, targeted AAVs at titers comparable to conventional production. This simple and generalizable synthetic-biology approach provides a versatile toolkit for rational AAV tropism engineering, offering a scalable route to custom vector design for research and therapeutic applications.

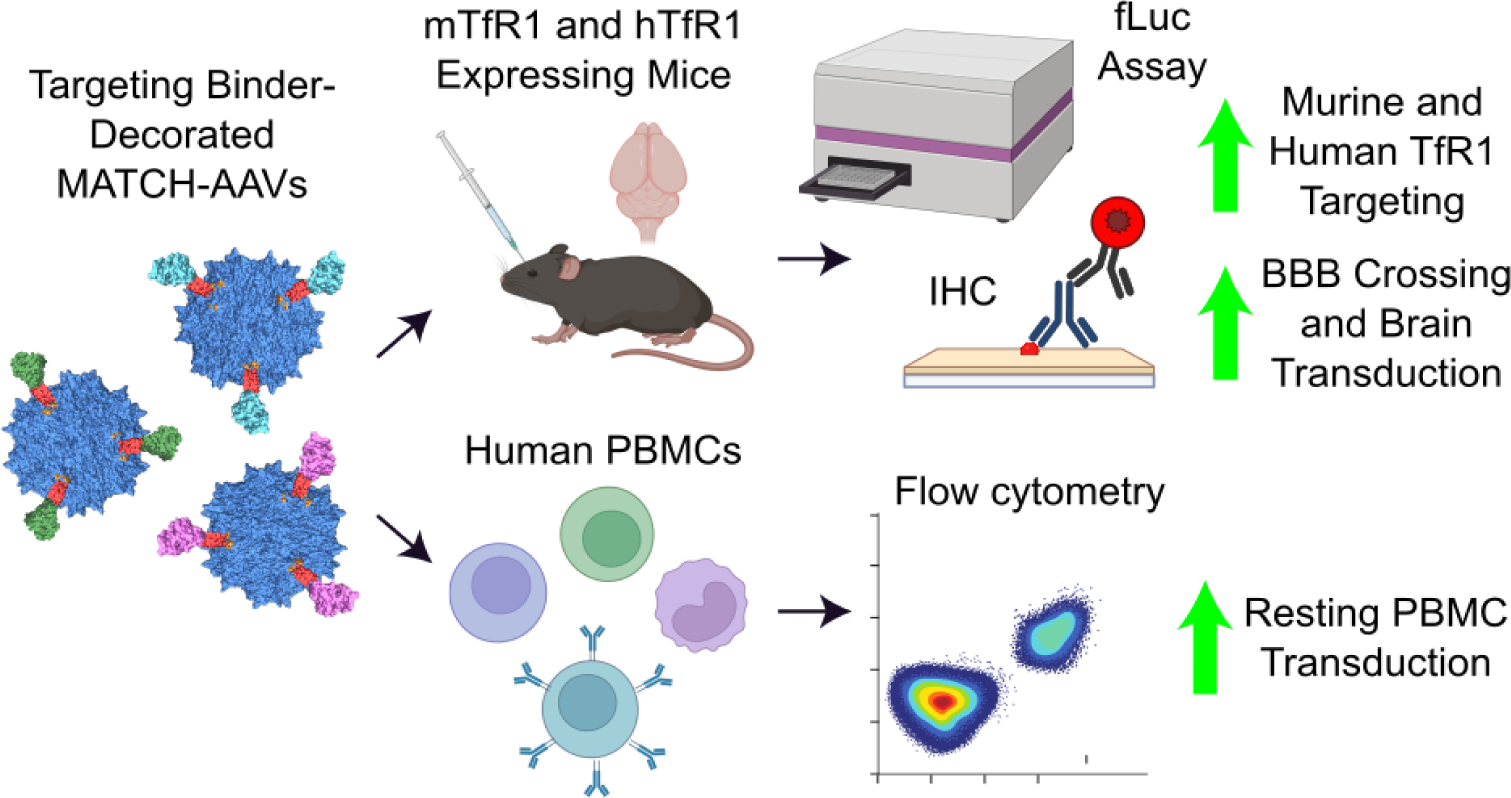

## Introduction

Gene therapy holds substantial promise for the treatment of numerous monogenic disorders^1^. Recombinant adeno-associated viruses (AAVs) remain the leading in vivo delivery platform due to their low immunogenicity, efficient transduction, and minimal genome integration^2^. However, despite several AAV-based therapeutics achieving regulatory approval, their clinical use is often constrained by dose-limiting toxicities^3^. Because natural serotypes display inefficient targeting of many therapeutically relevant cell types, current AAV therapies rely on high systemic vector doses, which can induce hepatotoxicity and provoke harmful immune responses^4,5^. The development of efficient, cell-type- and tissue-specific AAV vectors is therefore essential to improving the safety and efficacy of gene delivery.

To address the limitations of native AAV tropism, numerous studies have employed large-scale mutagenesis and peptide-insertion screens to engineer variants with enhanced specificity for muscle^6,7^, the central nervous system (CNS)^8,9^, liver^10,11^, and murine T lymphocytes^12^. While successful, these methods demand extensive resources and are restricted by the limited tolerance of the AAV capsid to lengthy peptide insertions^13,14^. To overcome these challenges, we and others have shown that larger biomolecules can be tethered onto the AAV capsid via incorporation of unnatural amino acids, enabling conjugation of diverse targeting motifs^15,16^. In parallel, recent efforts from our lab have leveraged natural ligand–receptor interactions by inserting short, ligand-derived peptide sequences into exposed capsid loops, enabling the identification of AAV variants with tissue-specific tropism^17^. Building on this experience, and inspired by work showing that the display of full-length targeting proteins on enveloped delivery vehicles can dramatically redirect tropism^18–20^, we sought to extend these principles to AAV through the conjugation of full-length proteins onto the exterior of the capsid.

Here, we introduce MATCH (Modulation of AAV Tropism through Conjugation to Homing proteins), a modular, scalable platform that enables efficient and versatile retargeting of AAV tropism through covalent conjugation of full-length targeting proteins. Building on our prior work^21^, MATCH involves the incorporation of a SpyTag peptide into the AAV capsid, and utilizes the SpyTag/SpyCatcher system, a peptide-protein pair derived from *Streptococcus pyogenes* that forms an irreversible covalent bond, enabling straightforward attachment of targeting motifs. By engineering a SpyTag motif directly into an exposed capsid loop, we create a recombinant AAV scaffold that can be rapidly functionalized with SpyCatcher-linked scFvs or proteins of interest. Using this platform, we identify a CD3-specific scFv that enables one-step activation and transduction of resting human T cells, and demonstrate that conjugation of transferrin receptor 1 (TfR1)-binding scFvs yields vectors capable of robust, TfR1-mediated CNS transduction in vivo, surpassing AAV9 performance in mice and achieving transduction efficiencies comparable to human TfR1-targeted AAV BI-hTfR1^22^ in human cells in vitro and humanized mice in vivo. Collectively, these findings establish MATCH as a flexible, clinically relevant strategy that builds directly on our prior work to overcome longstanding challenges in precise and programmable AAV tropism.

## Results

### Integration of SpyTag into the AAV capsid enables conjugation to SpyCatcher-linked homing proteins and receptor-targeted transduction

We sought to incorporate the SpyTag peptide directly into the AAV capsid to enable production of SpyTag-AAV using standard manufacturing methods. To do this, we inserted SpyTag001, a compact 13-amino acid variant compatible with the highly reactive SpyCatcher003^23^, into the primary receptor-binding domain of the AAV-DJ VP1 protein^24^ **(Fig. 1a)**. The SpyTag motif, flanked by flexible glycine–serine–serine (GSS) linkers, was positioned after residue 589 to generate AAV-DJ-SpyTag-VP, ensuring surface display of SpyTag on all capsid proteins (VP1/2/3) **(Supp. Fig. 1a)**.

**Figure 1.**
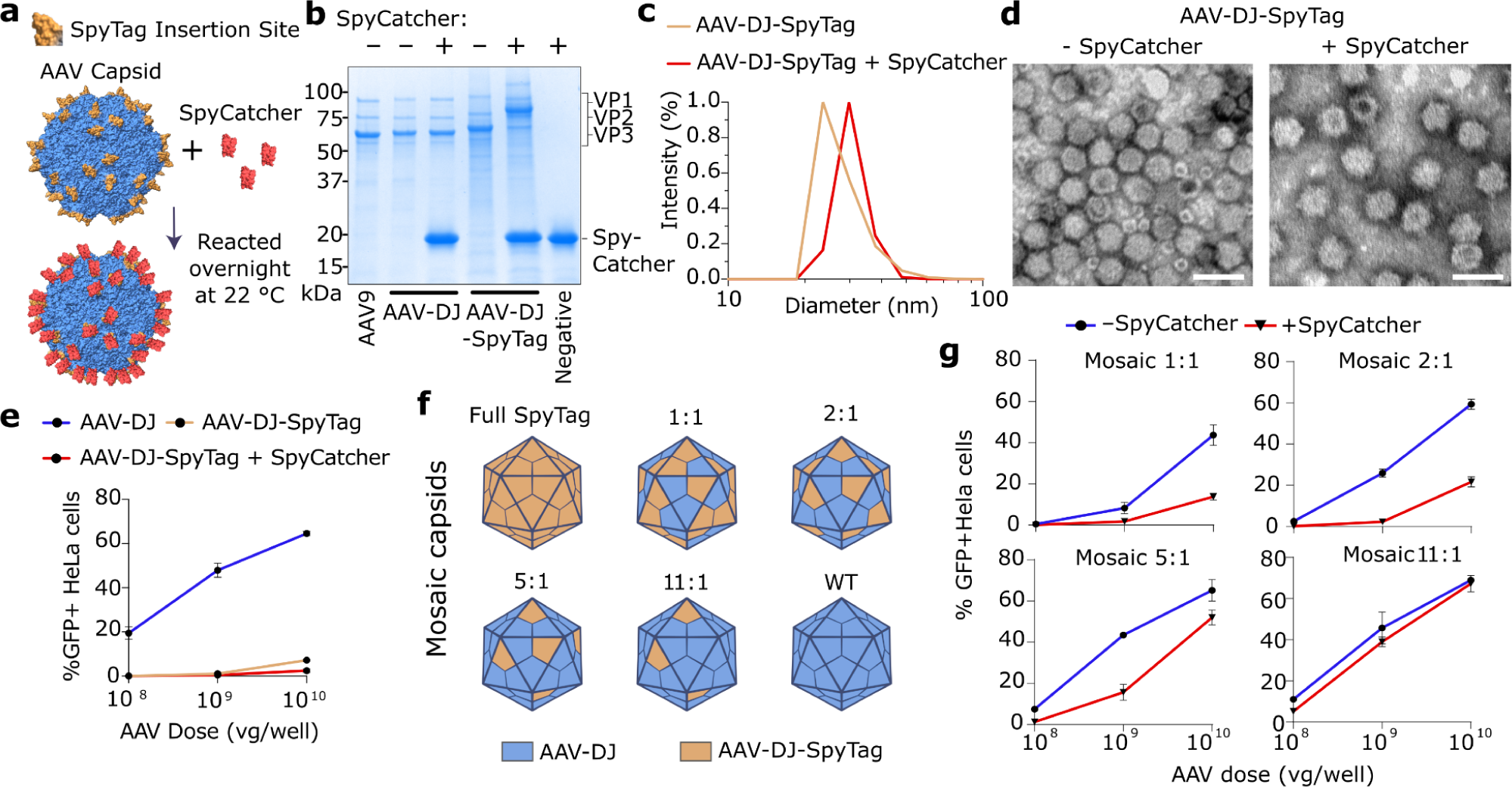
SpyTag insertion on the AAV capsid surface enables SpyCatcher conjugation. **(a)** Schematic of SpyTag insertion sites within the AAV-DJ capsid and conjugation to SpyCatcher. **(b)** Coomassie-stained gel showing VP1, VP2, and VP3 from 4.8 ✕ 10^11^ vg AAV9, AAV-DJ, and AAV-DJ-SpyTag ± incubation with 4.6 µg SpyCatcher. **(c)** Dynamic light scattering measurement of particle diameter and **(d)** transmission electron microscopy images of AAV-DJ-SpyTag capsids ± SpyCatcher. Scale bar: 50 nm. **(e)** Transduction efficiency of AAV-DJ and AAV-DJ-SpyTag in HeLa cells, with or without SpyCatcher. **(f)** Schematic of mosaic AAV capsids composed of AAV-DJ-VP and AAV-DJ-SpyTag-VP. **(g)** Transduction of HeLa cells with mosaic SpyTag-AAV-DJ capsids.

To confirm that SpyTag integration preserved reactivity to SpyCatcher, we produced AAV-DJ-SpyTag vectors packaging a GFP transgene, reacted them with recombinant SpyCatcher, and analyzed capsid proteins by SDS–PAGE **(Fig. 1b)**. All three viral proteins exhibited an upward shift relative to their AAV-DJ counterparts, consistent with incorporation of the 1.9 kDa SpyTag with GSS linkers. Following incubation with the 15.6 kDa SpyCatcher protein, a further shift was observed exclusively in the SpyTag-containing capsids, indicating successful SpyTag/SpyCatcher conjugation. Dynamic light scattering and transmission electron microscopy corroborated these findings by revealing increased particle size and altered capsid topography, respectively **(Fig. 1c,d)**. Notably, AAV-DJ-SpyTag exhibited a near-complete loss of transduction in HeLa cells both with and without SpyCatcher, demonstrating that full SpyTag occupancy disrupts native AAV-DJ receptor engagement **(Fig. 1e)**.

To restore productive transduction, we generated mosaic capsids composed of defined ratios of AAV-DJ-SpyTag-VP and AAV-DJ capsid proteins. Given the tripartite symmetry of the AAV protrusions, we co-transfected capsid plasmids at AAV-DJ:AAV-DJ-SpyTag ratios of 1:1, 2:1, 5:1, and 11:1, corresponding to mosaic capsids with an estimated 1.5, 1, 0.5, or 0.25 SpyTag motifs per protrusion **(Fig. 1f)**. Notably, all transfection ratios tested, including fully SpyTag capsids, produced titers comparable to the normal AAV-DJ capsid when packaging a GFP/Cre or fLuc transgene **(Supp. Fig. 1b,c).** Increasing incorporation of wild-type VP progressively restored transduction, both with and without bound SpyCatcher **(Fig. 1g)**. Additionally, nanoDSF analysis indicated that SpyTag VP incorporation did not reduce thermal stability of the 5:1 and 11:1 mosaic AAV capsids **(Supp. Fig. 1d)**. Based on these profiles, we selected the 5:1 and 11:1 formulations (MATCH-AAV-DJ 5:1 and 11:1) for subsequent studies.

### Conjugation of targeting ligands to mosaic AAV-DJ capsids modulates transduction efficiency and redirects tropism

Having established that mosaic SpyTag-containing capsids retain transduction competence, we next evaluated whether conjugation of SpyCatcher-linked ligands could redirect AAV tropism. We focused on two biologically and clinically relevant applications: transduction of resting human T cells ex vivo and CNS transduction in vivo.

Primary human CD3⁺ T lymphocytes, which include both CD4⁺ and CD8⁺ T cell populations and represent a major component of peripheral blood mononuclear cells (PBMCs), remain a challenging target for AAV-mediated gene delivery. While several AAV vectors engineered for T cell targeting have recently been developed, these systems typically leverage prior CD3/CD28-mediated activation to enhance transduction^25–27^. Additionally, adenoviral platforms have utilized IL-2 and CD3/CD28 binders to simultaneously activate and transduce T cells^28^. Recently, Cas9-packaging enveloped delivery vehicles (Cas9-EDVs) displaying CD3- and CD28-specific scFvs enabled simultaneous activation and transduction of resting T cells^19^; however, because this system relies on lentiviral components that integrate semi-randomly into the host genome, it raises concerns regarding insertional mutagenesis when delivering therapeutic transgenes. Thus, an AAV platform capable of activating and transducing T cells in a single step would be highly advantageous as a non-integrating alternative for applications such as CAR-T manufacturing and immune modulation^25,29^. Given the poor permissiveness of resting T cells to AAV, we screened a panel of SpyCatcher-fused scFvs targeting CD3 or CD28 for their ability to enhance MATCH-AAV-DJ transduction of human immune cells. Conjugation of these ligands to MATCH-AAV-DJ 5:1 and 11:1 identified αCD3-scFv-1-SpyCatcher as the most effective construct in a preliminary low-MOI screen **(Supp. Fig. 1e)**. When bound to MATCH-AAV-DJ 11:1, and purified using an iodixanol gradient to remove excess ligand, αCD3-scFv-1 conferred significant increase in transduction of resting human PBMCs compared to unmodified AAV-DJ, enabling transduction of ∼5% of a cell population that is otherwise minimally permissive to this capsid, representing an 84-fold increase in transduction over WT AAV-DJ **(Fig. 2a,b)**.

**Figure 2.**
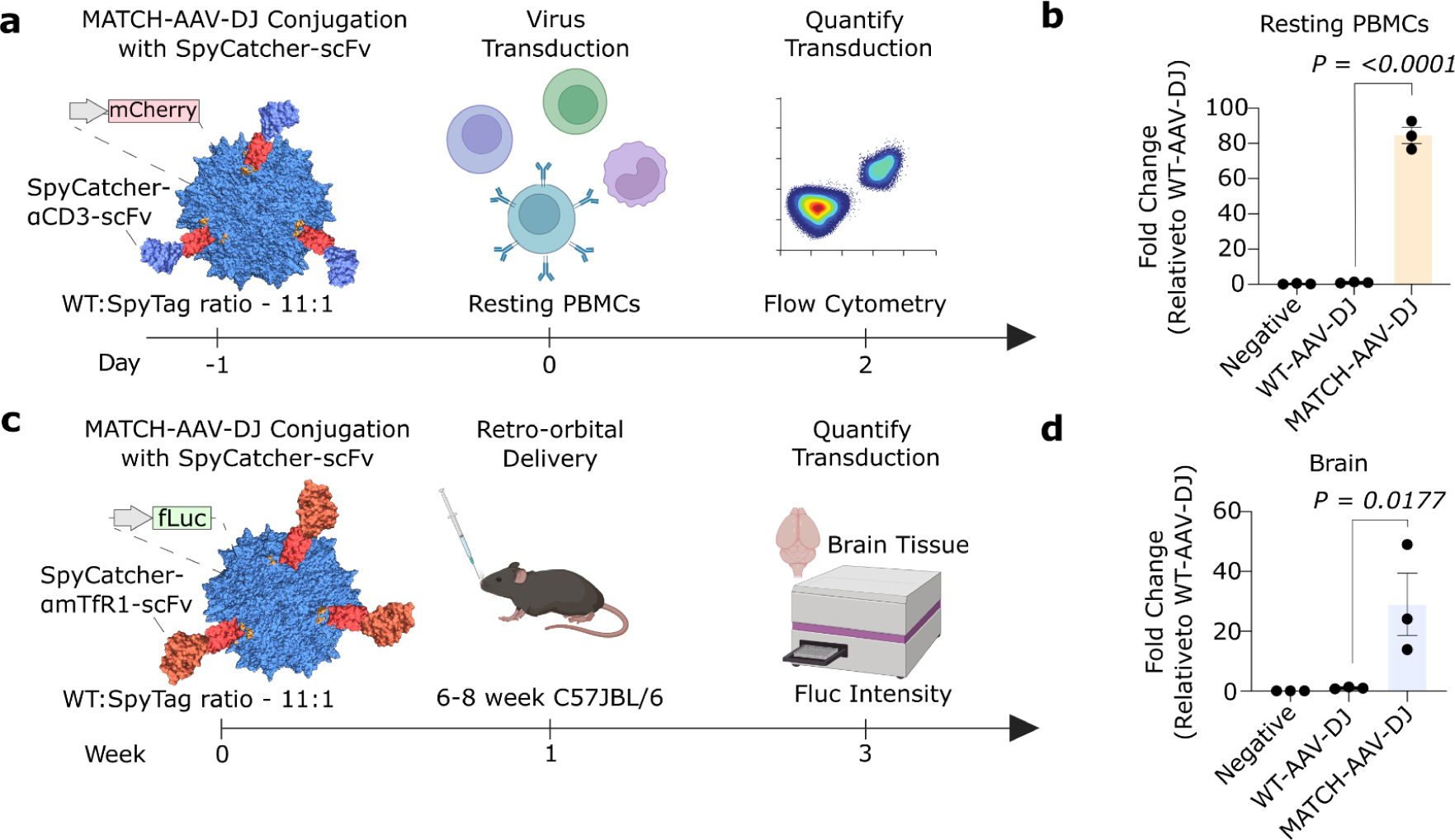
MATCH enables conjugation of targeting proteins to AAV-DJ and redirects tropism toward immune and CNS targets. **(a)** Workflow for transduction of human PBMCs using MATCH-AAV-DJ 11:1 conjugated to αCD3-scFv-1-SpyCatcher. **(b)** Fold increase in PBMC transduction relative to WT AAV-DJ when transduced at an MOI of 5 ✕ 10^5^ vg/cell, as quantified by flow cytometry 48 h post-transduction. **(c)** Workflow for in vivo CNS transduction using MATCH-AAV-DJ 11:1 conjugated to αmTfR1-scFv-SpyCatcher. **(d)** Fold increase in luciferase expression in mouse brain relative to AAV-DJ, three weeks after retro-orbital injection of 1 ✕ 10^12^ vg/mouse. Data represent mean ± SEM. One-way ANOVA with Tukey’s multiple comparisons test **(b,d)**.

The CNS represents another high-value target for gene therapy, where precise and efficient delivery has the potential to address numerous neurological disorders^30^. Although AAV9 can cross the blood–brain barrier^31,32^ and forms the basis of an FDA-approved therapy for spinal muscular atrophy^33^, severe dose-dependent toxicities including reported fatalities^3^ and prohibitive treatment costs^34^ underscore the need for more efficient, brain-targeted vectors. Transferrin receptor 1 (TfR1) is an attractive target due to its high expression on brain endothelial cells and its established ability to transport scFv-linked protein therapeutics across the blood-brain barrier (BBB)^35–38^. Recent work demonstrated that BI-hTfR1, an hTfR1-binding AAV9 variant generated through randomized peptide-insertion screening, achieves enhanced CNS transduction in hTfR1-expressing mice^22^, highlighting the potential of TfR1 engagement for AAV delivery. Motivated by these observations, we conjugated the high-affinity murine TfR1-binding scFv 8D3^36^, as well as a lower-affinity S101A variant^37^, to MATCH-AAV-DJ 5:1 and 11:1 capsids containing a firefly luciferase (fLuc) transgene. Following retro-orbital administration in mice and three weeks of expression, a bioluminescence analysis of brain and liver tissue revealed substantially elevated luciferase activity in the brain for both αmTfR1-scFv-conjugated vectors relative to AAV-DJ **(Fig. 2c,d; Supp. Fig. 1f,g)**.

Together, these results demonstrate that SpyCatcher-mediated conjugation of targeting scFvs effectively reprograms MATCH-AAV-DJ tropism, enabling enhanced gene delivery to both resting human T cells and the murine CNS.

### CD3-targeting epitopes conjugated to mosaic AAV9 capsids enable efficient activation and transduction of human PBMCs

To broaden the utility of MATCH, we adapted the SpyTag integration strategy to AAV9. SpyTag001 was introduced into two exposed capsid loops, Loop 1 (Q456–N457) and Loop 2 (A587–Q588), each flanked with either one or two GSS linkers to generate four AAV9-SpyTag-VP variants **(Fig. 3a)**. The two variants containing 2×GSS linkers were also incorporated into MATCH-AAV9 mosaics at an 11:1 WT:SpyTag ratio, yielding MATCH-AAV9 L1 and MATCH-AAV9 L2. All constructs produced AAV9 particles at titers comparable to wild type, and three fully SpyTag-containing capsids demonstrated robust reactivity with SpyCatcher **(Fig. 3b,c)**. Additionally, a nanoDSF analysis of MATCH-AAV9 L1 and MATCH-AAV9 L2 revealed similar thermal stability to WT AAV9 **(Supp. Fig. 1h)**.

**Figure 3.**
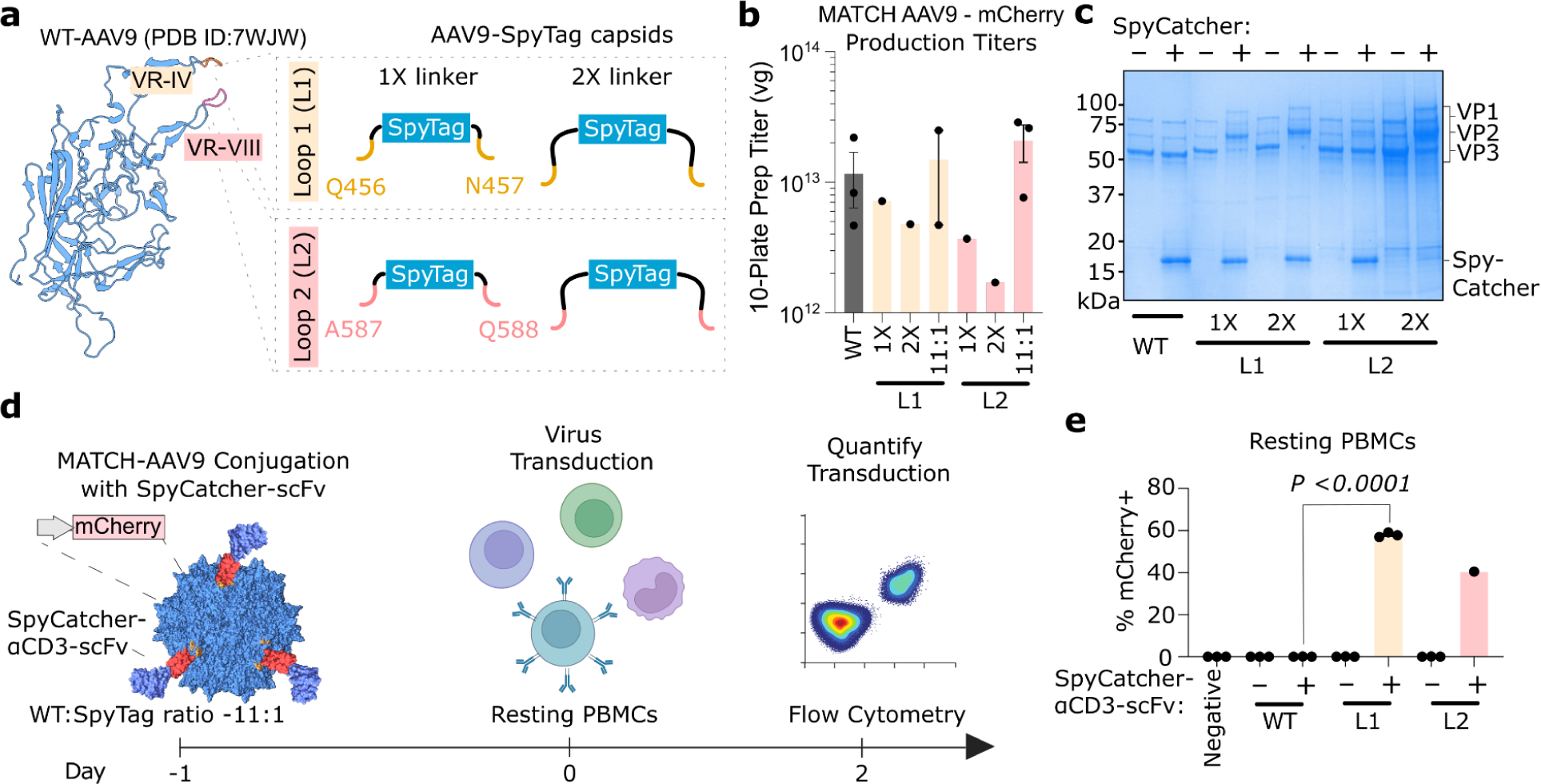
MATCH-AAV9 enables efficient transduction of resting human immune cells. **(a)** Schematic of AAV9 VP structure with SpyTag insertion sites, flanked by single or double GSS linkers. **(b)** Packaging efficiency of WT AAV9 and AAV9-SpyTag variants determined by qPCR titers against inverted terminal repeats. **(c)** Coomassie-stained viral protein profiles of 1 ✕ 10^11^ vg WT AAV9 and AAV9-SpyTag capsids ± incubation with 0.25 µg SpyCatcher. **(d)** Workflow for PBMC transduction using MATCH-AAV9 capsids conjugated to αCD3-scFv-1-SpyCatcher. **(e)** Percentage of transduced PBMCs measured by flow cytometry 48 h post-transduction at an MOI of 5 ✕ 10^5^ vg/cell. One-way ANOVA with Tukey’s multiple comparisons test **(e)**.

To determine whether MATCH-AAV9 recapitulated the enhanced immune-cell transduction observed with MATCH-AAV-DJ, we conjugated αCD3-scFv-1-SpyCatcher to each variant **(Fig. 3d)**. Both MATCH-AAV9 L1 and L2 efficiently transduced resting human PBMCs, with MATCH-AAV9 L1 transducing up to 58% of cells in vitro **(Fig. 3e)**. Transduction of PBMCs with αCD3-scFv-1-coated MATCH-AAV9 capsids encoding mCherry induced clear morphological hallmarks of T-cell activation, including cluster formation and increased cell size/granularity **(Supp. Fig. 2a,b)**. Following selection of live single cells **(Supp Fig. 2c,d)**, we observed that the vast majority of mCherry^+^ CD3^+^ T cells were also CD25⁺, indicating concurrent activation and transduction **(Supp. Fig. 2e,f)**.

### TfR1-targeted MATCH-AAV9 enables transferrin-receptor dependent transduction of the brain

We next evaluated whether MATCH-AAV9 could target the CNS through scFvs specific to the murine transferrin receptor (mTfR1). MATCH-AAV9 L1 and L2 were conjugated to αmTfR1-scFv-SpyCatcher and administered systemically via retro-orbital injection into mice. Three weeks post-injection, both variants produced substantially greater luciferase expression in the brain than WT AAV9 **(Fig. 4a,b, Supp. Fig. 3a)**. To further quantify vector delivery to the brain, we measured vector genomes and transgene expression by qPCR. MATCH-AAV vectors exhibited increased levels of transgenic DNA and mRNA, with transgenic mRNA levels in brain tissue comparable to those observed for the established murine brain transducer AAV-PHP.eB **(Supp. Fig. 3b,c,d)**^39^. Additionally, luciferase levels in the muscle and liver were assessed **(Supp. Fig. 3e,f)**. To determine whether brain transduction was restricted to the brain vasculature or distributed across the parenchyma, we performed confocal immunofluorescence (IF) analysis of brain sections. mCherry reporter expression was visualized along with CD31 to label endothelial cells, MAP2 to label neurons, and DAPI to mark nuclei. In C57BL/6 mice, WT AAV9 transduction was largely confined to CD31⁺ endothelium. In contrast, MATCH-AAV vectors targeted to mTfR1 through conjugation to 8D3 exhibited reporter expression outside CD31⁺ vasculature, with transgene appearing throughout the tissue **(Supp. Fig. 4)**. These orthogonal measurements complement the luciferase activity data and support enhanced brain delivery by TfR1-targeted MATCH-AAV constructs.

**Figure 4.**
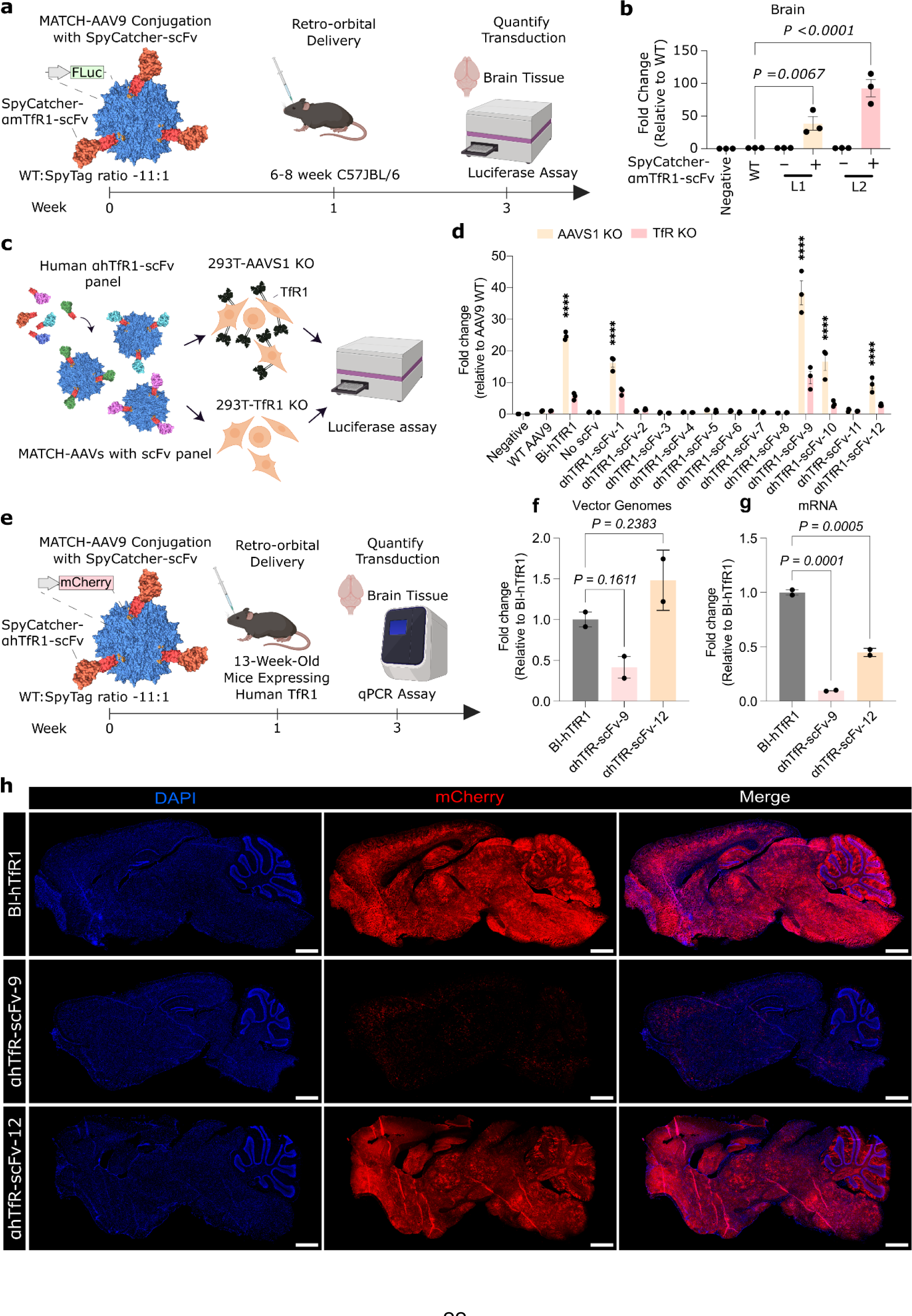
MATCH-AAV9 enables efficient transduction of the central nervous system. **(a)** Workflow for in vivo CNS transduction using MATCH-AAV9 conjugated to αmTfR1-scFv-SpyCatcher. **(b)** Fold improvement in brain luciferase expression over WT AAV9, three weeks after retro-orbital injection of 1 ✕ 10^12^ vg/mouse. One-way ANOVA with Tukey’s multiple comparisons test was used with WT AAV9 as the main comparison group. **(c)** Workflow for screening αhTfR1-scFv-SpyCatcher variants in AAVS1 KO and TfR1 KO HEK293T cells. **(d)** Fold improvement in luciferase expression over WT AAV9 in AAVS1 KO and TfR1 KO cells 48 h post-transduction using MATCH-AAV9 L2 decorated with αhTfR1-scFv-SpyCatcher variants at an MOI of 1 ✕ 10^4^ vg/cell. Data represent mean ± SEM. ****P < 0.0001. Two-way ANOVA with Dunnett’s multiple comparisons test was used with WT AAV9 as the main comparison group. **(e)** Workflow for in vivo CNS transduction using MATCH-AAV9-L2 conjugated to αhTfR1-scFv-9 or αhTfR1-scFv-12. Prevalence of MATCH-AAV9 **(f)** vector genomes and **(g)** mRNA transcripts in the murine brain relative to BI-hTfR1, three weeks after retro-orbital injection of 1 ✕ 10^12^ vg/mouse as determined by qPCR. One-way ANOVA with Tukey’s multiple comparisons test was used with BI-hTfR1 as the main comparison group. **(h)** Sagittal brain sections immunostained for nuclei (DAPI) and AAV transgene (mCherry) from mice three weeks after retro-orbital delivery of 1 ✕ 10^12^ vg/mouse BI-hTfR1 or MATCH-AAV9 conjugated to αhTfR1-scFv-9 or αhTfR1-scFv-12. Scale bar 1mm.

To assess targeting of human TfR1, we generated a panel of scFvs against hTfR1 and conjugated them to MATCH-AAV9 L2. WT AAV9 and BI-hTfR1, a previously engineered hTfR1-binding variant, served as comparators. These vectors were used to transduce HEK293T cells in which either the AAVS1 locus or hTfR1 was knocked out via CRISPR–Cas9 **(Fig. 4c)**. The AAVS1 knockout line served as a CRISPR-edited control, as disruption of this neutral locus is not expected to affect AAV entry or TfR1 expression. Several hTfR1-binding scFvs markedly enhanced MATCH-AAV9 L2 transduction in AAVS1-knockout cells, achieving efficiencies comparable to BI-hTfR1 **(Fig. 4d)**. This enhancement was diminished in TfR1-knockout cells, supporting a receptor-dependent mechanism of transduction. Similar trends were observed in wild-type HEK293T cells **(Supp. Fig. 3a,b)**.

We next evaluated brain delivery in vivo using humanized B-hTfR1 mice expressing the apical domain of hTfR1. Quantification of vector genomes and transgene mRNA in brain tissue revealed that MATCH-AAV vectors conjugated to αhTfR-scFv-12 achieved levels of transgene delivery and transcription comparable to BI-hTfR1 **(Fig. 4f,g)**. Immunohistochemical analysis of mCherry reporter expression further demonstrated widespread signal across multiple brain regions following systemic administration **(Fig. 4h; Supp. Fig. 5c,d)**. This expression was also observed outside CD31⁺ endothelial structures and broadly distributed through the parenchyma, indicating efficient BBB-crossing **(Supp. Fig. 6,7)**.

Following the identification of efficient T cell and CNS targeted vectors, we elected to improve the efficiency and broader applicability of the MATCH system by streamlining vector production. While separate production of MATCH capsids and targeting ligands enables modular retargeting, a one-pot transfection approach would simplify vector generation and eliminate additional purification steps. To investigate this, we co-transfected targeting plasmids encoding either αCD3-scFv-1-SpyCatcher or αhTfR1-scFv-12-SpyCatcher alongside the standard triple-transfection MATCH-AAV9 L2 plasmid pool (11:1 WT:SpyTag Rep/Cap, pAd-helper, and transgene) **(Fig. 5a)**. Targeting plasmids were included at mass ratios of 1:3 or 1:7 relative to the MATCH plasmids. Vectors carried either an mCherry (CD3-targeted) or fLuc (hTfR1-targeted) transgene. These “Mix-and-MATCH” productions yielded titers comparable to WT AAV9 using our standard AAV production protocol **(Fig. 5b)**. Upon application to resting human PBMCs, CD3-targeted Mix-and-MATCH AAV9 vectors transduced a significant portion of the T cell population **(Fig. 5c,d)**. Additionally, hTfR1-targeted vectors displayed significantly enhanced transduction of HEK cells in comparison to WT AAV9 **(Fig. 5e,f)**. Collectively, these data indicate that one-pot “Mix-and-MATCH” productions maintain ligand-dependent targeting and transduction efficiency while streamlining MATCH vector manufacturing in a manner analogous to standard AAV production workflows.

**Figure 5.**
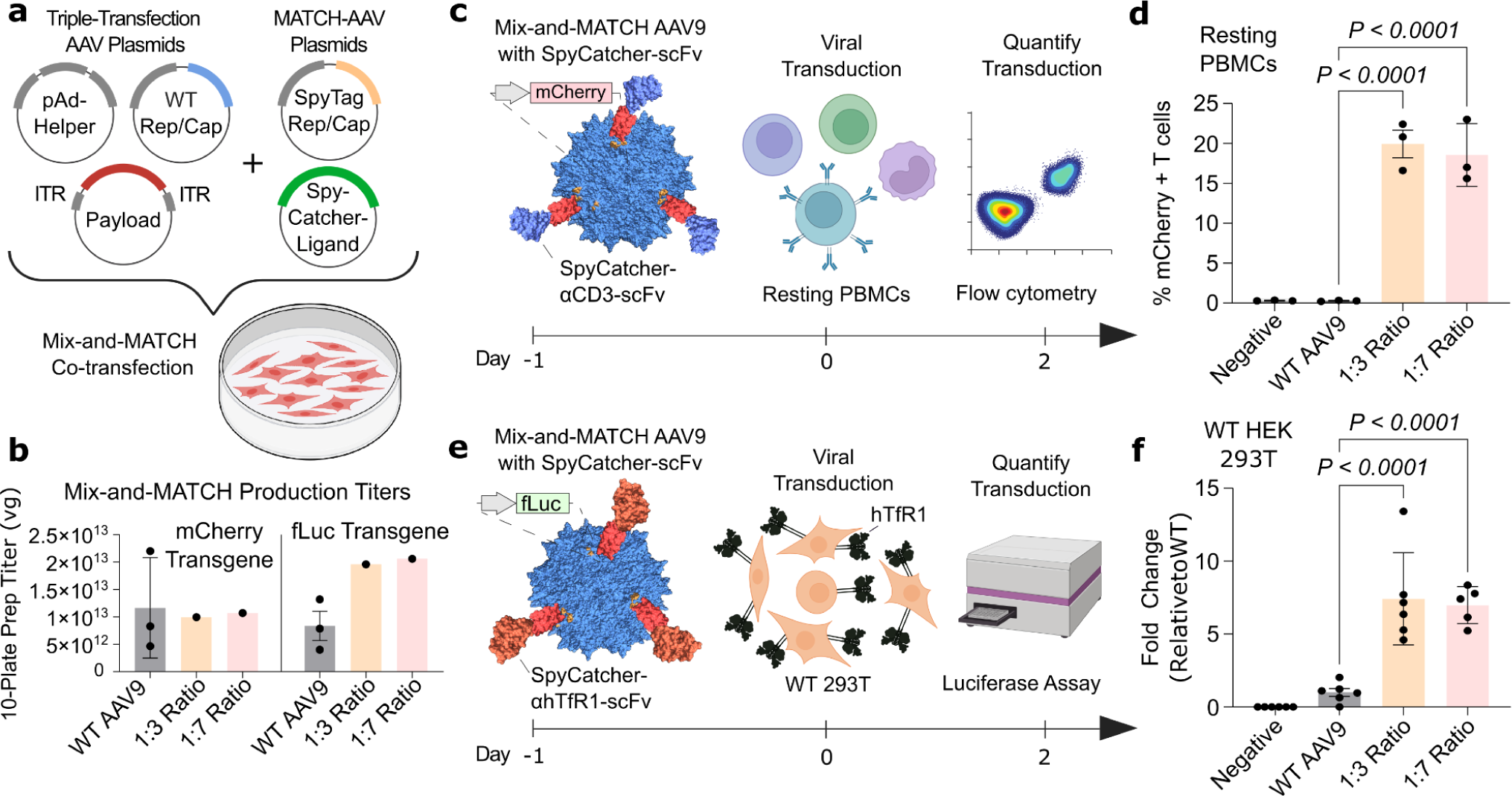
Joint transfection of MATCH AAV9 L2 and targeting scFvs enables one-pot production of MATCH vectors. **(a)** Workflow for one-pot Mix-and-MATCH AAV production. **(b)** Packaging efficiency of Mix-and-MATCH AAV9 variants in comparison to WT AAV9 as determined by qPCR using primers against inverted terminal repeats. **(c)** Workflow for PBMC transduction using Mix-and-MATCH-AAV9 L2 capsids co-transfected with αCD3-scFv-1-SpyCatcher. **(d)** Percentage of transduced T cells as measured by flow cytometry 48 h post-transduction at an MOI of 5 ✕ 10^5^ vg/cell. **(e)** Workflow for transduction of WT HEK293T cells with Mix-and-MATCH-AAV9 L2 capsids co-transfected with αhTfR1-scFv-SpyCatcher. **(f)** Relative fLuc signal intensity from HEK293T cells transduced with Mix-and-MATCH AAV9 variants at an MOI of 1 ✕ 10^4^ vg/cell in comparison to WT AAV. One-way ANOVA with Tukey’s multiple comparisons test **(c, f)**.

## Discussion

In this study, we present MATCH, a modular approach to AAV engineering that addresses several longstanding challenges in vector development. Traditional AAV discovery strategies rely heavily on peptide-insertion mutagenesis, yet the AAV capsid can tolerate only short peptides at a limited number of sites^13,14^, restricting the incorporation of larger or structurally complex receptor-binding motifs. MATCH circumvents these constraints by enabling the conjugation of full-sized targeting proteins directly onto the exterior of the capsid. Through sparse incorporation of SpyTag on the capsid surface and subsequent covalent linkage to SpyCatcher-fused scFvs, we demonstrate efficient transduction of resting PBMCs in vitro and robust targeting of the murine CNS in vivo. Importantly, these effects were observed across two widely used parental serotypes, AAV-DJ and AAV9, with MATCH vectors outperforming their unmodified counterparts in each application.

Previous methods for modular AAV retargeting rely on unnatural amino acid incorporation into the AAV capsid, followed by click-chemistry based linkage directly to a targeting motif^15^ or peptide^40^. Enzymatic capsid labeling strategies have also been explored, including insertion of a biotin acceptor peptide into the capsid followed by BirA-mediated ligation of functional probes or targeting ligands^41^. In the MATCH system, SpyTag containing capsids are generated using standard AAV production methods, eliminating the need for genetic code expansion and enzymatic processing. Other studies have incorporated large targeting ligands, such as DARPins, directly into capsid proteins, though this necessitates either additional purification steps to enrich functional particles^42^, or careful optimization of insertion site and stoichiometry to ensure capsid assembly and gene transfer^26^. In contrast, the MATCH system enables the targeting of a common capsid to multiple biologically distinct targets without the need for additional purification or extensive optimization.

Several recent and concurrent studies have also reported SpyTag/SpyCatcher-based approaches for modular AAV retargeting. These systems either incorporate SpyTag on every protein of the viral capsid or restrict it to specific VP components, resulting in vectors with fixed ligand valency defined during capsid assembly^43,44^. In contrast, the MATCH platform employs mosaic capsids generated through controlled mixing of wild-type and SpyTag-containing capsid plasmids, allowing AAVR engagement and enabling tunable ligand display across multiple incorporation ratios and capsid backgrounds. Notably, whereas prior studies have primarily focused on tumor-targeting applications^43–45^ MATCH demonstrates retargeting in biologically distinct contexts, including enhanced delivery to the brain, a major target of clinical AAV gene therapy, and efficient transduction of primary resting T cells.

Our findings also highlight key structural and functional considerations for SpyTag-modified AAVs. Specifically, SpyTag insertion in AAV-DJ was positioned immediately after Asn589, a residue located within the heparan sulfate proteoglycan (HSPG) binding domain defined by Arg587 and Arg590^24^. The marked reduction in HeLa cell transduction observed with fully SpyTag-modified capsids, or even with mosaic capsids carrying a substantial SpyTag fraction, is therefore consistent with disruption of HSPG engagement. Notably, the additional decline in transduction following SpyCatcher binding suggests steric interference with secondary receptor interactions as well. AAVR, an essential secondary receptor for multiple AAV serotypes including AAV2, AAV8, and AAV9, the parents of AAV-DJ, binds in close proximity to the threefold axis where SpyTag was inserted^46–49^. Thus, conjugated SpyCatcher likely impedes AAVR engagement, further limiting internalization and endosomal trafficking. Incorporating SpyTag into only a minority of capsid subunits, as implemented in MATCH, preserves access to native receptor pathways while simultaneously enabling new receptor interactions through conjugated ligands. Consistent with this model, the parent serotype influences how MATCH vectors integrate native and engineered targeting cues. AAV9 uses galactose as its primary receptor, binding through residues that form a pocket at the base of the threefold protrusion in a region distinct from our SpyTag insertion sites^50^. By deriving MATCH-AAVs from parent serotypes with different receptor dependencies, it may be possible to combine or orthogonally tune multiple receptor interactions, providing enhanced control over cell-type specificity and tissue targeting.

Our work identifies CD3 as a particularly powerful targeting receptor for T-cell gene delivery. CD3 is a central component of the T-cell receptor complex and mediates activation signaling upon antigen engagement^51,52^. Following stimulation, CD3 undergoes rapid internalization and recycling or degradation^53,54^. Antibody- or scFv-mediated CD3 engagement can recapitulate this process, leading to T-cell activation^55^. Accordingly, the benefits of targeting CD3 with MATCH likely arise from three synergistic mechanisms: (1) high-affinity attachment and concentration of AAV on the T-cell surface; (2) CD3-triggered internalization that co-traffics the bound vector into the cell; and (3) simultaneous activation of the T cell, which may enhance its susceptibility to AAV transduction and facilitate downstream therapeutic programming. Notably, MATCH-AAV enabled robust transduction and activation of resting human T cells, an outcome not previously achieved by AAV vectors without prior T-cell activation.

Similarly, conjugation of an established anti-mTfR1 scFv (8D3) permitted highly efficient CNS targeting in vivo using both MATCH-AAV-DJ and MATCH-AAV9. The extent of brain transduction observed far exceeded that achieved by the unmodified parental serotypes. In parallel, screening a panel of hTfR1-targeting scFvs revealed several binders that achieved transduction efficiencies comparable to BI-hTfR1 in human cells. The substantial attenuation of transduction in hTfR1 knockout cells, but not in AAVS1 knockout controls, confirms that MATCH-mediated enhancement is hTfR1-dependent. MATCH-AAV vectors conjugated to mTfR1- and hTfR1-binding scFvs mediated transduction across brain regions and throughout the parenchyma, with widespread reporter expression outside CD31⁺ endothelial structures. These findings demonstrate that MATCH can be used to effectively engage clinically relevant human receptors, and support its translational potential for CNS gene delivery.

Interestingly, we observed substantial variability in AAV transduction efficiency among the tested anti-TfR scFvs. These scFvs were derived from reported heavy and light chain sequences of antibodies known to bind the human transferrin receptor as reported by the PLAbDab database (αhTfR1-scFvs 1-10)^56^ and the literature (αhTfR1-scFvs 11-12)^57,58^, which were reformatted as scFv constructs for compatibility with the MATCH system. Among these candidates, an scFv derived from the CH3 region of TV35 (PDB ID: 6W3H)^58^, αhTfR1-scFv-12, showed the most robust functional activity. Prior studies on antibody-mediated targeting of TfR for blood–brain barrier delivery have demonstrated that relatively small differences in antibody affinity, epitope location, or receptor trafficking behavior can strongly influence transport and intracellular routing outcomes^37,38,59^. It is therefore plausible that the differential performance observed here reflects differences in the biophysical nature of the scFv–TfR interaction that influence receptor engagement and subsequent viral entry. While a systematic analysis of these parameters was beyond the scope of the present work, the modular nature of the MATCH platform provides a useful framework for rapidly screening such ligand properties in future studies.

A key strength of the MATCH system is the simplicity with which vectors of diverse tropisms can be generated. Parent capsids require only sparse SpyTag incorporation, a modification that does not significantly impair production yield or stability, and targeting proteins can be produced, purified, and stored independently before conjugation. This modularity supports scalable, on-demand vector assembly, enabling the same core capsid and transgene cassette to be readily redirected to distinct cell types by swapping targeting ligands. It should be noted that, while mosaic capsid assembly enables tunable ligand display, individual particles likely exhibit a distribution of SpyTag incorporation which may introduce heterogeneity in ligand density at the single-capsid level. As an alternative to the modular MATCH system, we also developed the one-pot Mix-and-MATCH productions approach to enable generation of targeted vectors using standard AAV production techniques, analogous to standard triple-transfection protocols.

In summary, our results demonstrate that MATCH enables efficient and programmable AAV targeting through the one-step conjugation of full-length proteins to a shared capsid framework. As the MATCH system develops, we anticipate that it will provide a versatile platform for addressing a broad spectrum of gene delivery challenges across diverse tissues and cell types, and facilitate the identification of useful receptors and ligands for targeted gene therapies.

## ACKNOWLEDGEMENTS

The authors thank Dhruva Katrekar, Dr. Sami Nourreddine, and Duy An Le. Figures were created in part using BioRender, and statistical analyses and data visualization were performed using GraphPad Prism. Our work is generously supported by UCSD Institutional Funds, NIH grants (R01HG012351, R01CA310063, R01NS131560), a CIRM Grant (DISC4-19271), a Department of Defense Grant (W81XWH-22-1-0401) and a UC San Diego Gene Therapy Initiative Grant (GTI-2024-018). The authors thank the University of California, San Diego - Cellular and Molecular Medicine Electron Microscopy Core (UCSD-CMM-EM Core, RRID: SCR_022039) for equipment access and technical assistance. The authors thank the Nikon Imaging Center at UCSD for their support with confocal microscopy experiments, and the UCSD SOM Microscopy Core (NS047101, OD030505, OD036455) for assisting with tissue section tilescanning. The authors thank the Stem Cell Genomics Core at the Sanford Stem Cell Institute for assistance with cell characterization via flow cytometry. The authors would also like to thank the microscopy core in the UCSD neurosciences department which is supported by a NIH grant (NINDS P30NS047101), the La Jolla Institute Histology Core facility for their expert help with tissue preparation and cryosectioning, and the GT3 Core Facility of the Salk Institute which is supported with funding from NIH-NCI CCSG: P30 014195, a NINDS R24 Core Grant, and funding from NEI.

## AUTHOR CONTRIBUTIONS

N.G. and P.M. conceived and planned the study. N.G., S.K., J.R., S.Y., A.P., and B.S. performed experiments. N.G. analyzed the data. N.G. and and S.K. generated figures. N.G., S.K., and P.M wrote the manuscript, and all authors assisted in editing the manuscript.

## DATA AVAILABILITY

The data that support the findings of this study are available from the corresponding author upon reasonable request.

## MATERIALS AND METHODS

### Cell culture and in vitro transductions

HEK293T cells (ATCC, CVCL_0063), HEK293T-derived cell lines, and HeLa cells (ATCC, CVCL_0030) were maintained in high glucose DMEM with GlutaMAX^TM^ (Gibco, 10566024), supplemented with 10% FBS (Gibco, A5256801) and 1% Penicillin-Streptomycin (Gibco, 15140122). HEK293T cells, HEK293T-derived cell lines, HeLa cells, and PBMCs were cultured in a 37 °C incubator with 5% CO2. FreeStyle^TM^ 293-F cells (ThermoFisher, R79007) were maintained in FreeStyle™ 293 Expression Medium (Gibco, 12338026) in a shaking 37 °C incubator with 8% CO2. AAVS1 (guide sequence: GGCGATATCTAGGTAGCCAC) and TfR1 (guide sequence: GAATACCTCTAGCCATTCAG) knockouts were conducted using lentiCRISPR v2 (Addgene #52961). All AAV transductions of HEK293T cells, HEK293T-derived cell lines, and HeLa cells were performed in maintenance media, with the vector-containing solution added consisting of under 20% of the total culture volume. For HeLa cell transduction experiments, 3 ✕ 10^4^ cells were seeded 24 hours before transduction, with positive cell counts determined via flow cytometry 48 hours post transduction. For HEK293T cell transduction experiments, cells were seeded 24 hours before transduction. 48 hours after transduction, media was changed gently and relative cell viability per well was determined using a CCK8 assay, with an incubation time of 1.5 hours (Dojindo, CK04-01) before luciferase activity was assessed using the Bright-Glo™ Luciferase Assay System (Promega, E2610) according to manufacturer’s protocols. Luciferase signal was then normalized to relative cell viability.

Human PBMCs were obtained from either Charles River Laboratories (PB009C-2) or STEMCELL Technologies (70025.2), and were maintained in RPMI 1640 (Gibco, 11875093) supplemented with 10% FBS and Human Recombinant 180 IU/mL IL-2, ACF (STEMCELL Technologies, 78145). All AAV transductions of immune cells were performed in culture media without FBS, with the vector-containing solution added consisting of under 10% of the total culture volume. FBS was added to 10% of the total culture volume two hours after transduction.

### Vector construction

SpyTag and SpyCatcher sequences were based on previous works from the lab of Mark Howarth. SpyTag001 flanked by Glycine-Serine-Serine (GSS) linkers were obtained as eBlocks^TM^ gene fragments for cloning into the AAV capsid loops. Recombinant transgene vectors were cloned into the AAV plasmid cloning backbone pZac2.1 (University of Pennsylvania Vector Core) flanking a CMV promoter using Gibson assembly. All SpyCatcher constructs were cloned into a pCAG backbone (Addgene Plasmid #11150). The coding sequence of SpyCatcher003 was codon optimized and fused at the C-terminus of ligands following a GGGGS linker. A Igκ leader sequence was included on the N-terminus of targeting ligands for secretion, and a 6X His-tag was added directly at the C-terminal end of SpyCatcher003 for purification by affinity chromatography. Therefore, each construct consisted of the following components in order from N-terminus to C-terminus: Igκ leader sequence, scFv, GGGGS, SpyCatcher, 6xHis-Tag.

### Protein production and purification

For individual production of SpyCatcher-linked proteins, 1 µg/mL of plasmid DNA was transfected into 293-F cells when cell density reached ∼1 ✕ 10^6^ cells/mL using PEI (Kyfora Bio, 23966-1) at a 1:1 DNA:PEI mass ratio. Cell suspension supernatants were harvested 5 days later, sterile-filtered (0.45 µm), and purified using affinity chromatography with HisTrap HP columns (Cytiva, 17524802) according to manufacturer protocols. Purified protein fractions were then concentrated using Amicon® 3K centrifugal filters (Merck, UFC800324) and washed three times in 1✕ PBS, pH 7.4 (Gibco, 10010-049) with 50 mM sodium chloride and 0.0001% Pluronic F68 (ThermoFisher, 24040-032). Protein titers were determined using Coomassie Blue staining with protein standards and stored at 4 °C until use.

### AAV production

AAV-PhP.eB (Addgene #103005) and BI-hTfR1 (Addgene #218796) sequences were obtained from Addgene, originally deposited by the laboratories of Viviana Gradinaru and Ben Deverman. AAVs were prepared using the triple transfection method followed by iodixanol gradient purification. Briefly, HEK293T cells were plated on 15cm plates 48 hours before transfection. Media was changed 2 hours before transfection. For all AAV transfections, 30 µg of DNA was added per 15cm plate, using PEI at a DNA:PEI mass ratio of 1:5 as previously described^15^. Standard AAV transfections were done with 10 µg each of transgene, pHelper and respective Rep/Cap plasmids per plate. For MATCH-AAV preps, the 10 µg of Rep/Cap plasmid was split between parental and Spy-Tag containing Rep/Cap plasmids at the designated ratio (1:1, 2:1, 5:1, or 11:1 parental:SpyTag-containing). For Mix-and-MATCH preps, the amount of the transgene, pHelper and respective Rep/Cap plasmids was reduced to 7.5 (1:3) or 8.75 (1:8) µg DNA per plasmid per plate, with SpyCatcher-scFv plasmid comprising the remaining 7.5 (1:3) or 3.75 (1:8) ug DNA per plate. Media was changed again 24 hours after transfection. Cells were harvested 84 hours after transfection and purified using iodixanol density gradient ultracentrifugation. Purified AAVs were then dialyzed and concentrated with AAV buffer (1 ✕ PBS, with 50 mM sodium chloride and 0.0001% Pluronic F68 (ThermoFisher, 24040-032)) using Amicon 50K centrifugal filters (Merck, UFC905024) to a final volume of less than 500 µL. Viral titers were determined using qPCR with primers specific to the ITRs and iTaq Universal SYBR Green Supermix (Bio-Rad, 1725122) for 40 cycles through comparison to a standard (ATCC VR-1616). AAVs were aliquotted and stored at -80 °C until use. DLS readings were taken using a Wyatt Instruments DynaPro Plate Reader III. Thermal stability measurements were performed using nano differential scanning fluorimetry (nanoDSF) on a Prometheus Panta instrument (NanoTemper Technologies), with intrinsic fluorescence monitored during a thermal ramp with data acquired at 1 °C increments.

### AAV-SpyTag and Ligand-SpyCatcher reactions

SpyCatcher003 for initial conjugation experiments was obtained from Kerafast (EOX003). AAV-SpyTag vectors were incubated with 100 molar excess of ligand-SpyCatcher molecules at room temperature overnight in AAV buffer. For all TfR-targeting in vitro experiments and the initial low-MOI CD3/CD28 scFv panel test, reactions were washed once with AAV buffer, concentrated to under 30 µL using Amicon 100K centrifugal filters (Merck, UFC510024), and titered using qPCR before application to cells. For all other PBMC transduction experiments, reactions were purified using iodixanol density gradient ultracentrifugation before being dialyzed and concentrated to a volume of less than 500 µL with AAV buffer using Amicon 50K centrifugal filters. This volume was concentrated further to under 30 µL using Amicon 100K centrifugal filters and titered using qPCR before experiments.

### Flow cytometry

For general transduction experiments, 48 hours after AAV transduction, cells were centrifuged and resuspended in 1✕ PBS with 2% FBS and 0.1 µg/mL DAPI Staining Solution (Miltenyi Biotec, 130-111-570), pH 7.4 at 1 ✕ 106 cells/mL and analyzed on the BD LSRFortessa™ Cell Analyzer, gating for live cells and mCherry or GFP fluorescence. Transduction efficiencies were determined by mCherry or GFP positive cells. Fold changes were determined by normalized transduction efficiencies to WT controls. For CD3 and CD25 labeling experiments, cells were centrifuged and resuspended in 1✕ PBS with 2% FBS, pH 7.4 at 1 ✕ 10^6^ cells/mL with Human TruStain FcX™ (BioLegend, 422301). After 10 minutes, cells were stained with APC anti-human CD3 Antibody (BioLegend, 300412) and FITC anti-human CD25 Antibody (BioLegend, 302603) for 30 minutes on ice. Cells were washed once with 1✕ PBS with 2% FBS, pH 7.4, and finally resuspended in 1✕ PBS with 2% FBS and 0.1 µg/mL DAPI Staining Solution (Miltenyi Biotec, 130-111-570), pH 7.4 and analyzed on the BD LSRFortessa™ Cell Analyzer.

### Animal experiments

All animal procedures were performed in accordance with protocol S16003 approved by the Institutional Animal Care and Use Committee of the University of California San Diego. C57BL/6J (Jackson Laboratories, 000664) or B-hTfR1 mice (Biocytogen, 110861) were housed at a temperature of ∼70°F (21°C), with ∼55% humidity and a 12-h light/dark cycle with food and water ad libitum.

### In vivo Luciferase assay

For in vivo experiments with a luciferase readout, 6-8 week old male C57BL/6J mice were injected retro-orbitally with 1 ✕ 10^12^ vector genomes per mouse and euthanized three weeks post AAV delivery, with 3 mice per treatment group. Whole brain, gastrocnemius muscle and left-lobe liver tissues were harvested, snap-frozen in liquid nitrogen and stored at -80 °C until use. Tissues were homogenized using bead mill tubes (Fisher Scientific, 15-340-154) in the TissueLyser LT (Qiagen, 85600) before luciferase detection with the Bright-Glo Luciferase Assay system (Promega, E2610) as per the manufacturer’s instructions. Luciferase activity was normalized to protein concentration using absorbance data from Pierce™ BCA Protein Assay Kits (Thermo Scientific, 23225).

### Tissue preparation and immunohistochemistry

For immunofluorescence in-vivo experiments comparing WT AAV9 and MATCH-AAV9 L2, 6-8 week old male C57BL/6J mice were injected retro-orbitally with 5 ✕ 10^12^ vector genomes per mouse, with 2 mice per treatment group. For immunofluorescence in-vivo experiments comparing BI-hTfR1 and MATCH-AAV9 L2 constructs, 13 week old male B-hTfR1 mice were injected retro-orbitally with 1 ✕ 10^12^ vector genomes per mouse, with 2 mice per treatment group. For all immunofluorescence in-vivo experiments, after three weeks, mice were anesthetized with ketamine hydrochloride (Ketaset^®^, NDC 54771-2013-1) and xylazine (AnaSed®, NDC 59399-110-20) at 100mg/kg and 11mg/kg of body weight respectively, and perfused with chilled 4% paraformaldehyde (PFA) in 1✕ PBS. Harvested tissues were then post-fixed overnight (C57BL/6J experiments) or for 5 hours (BI-hTfR1 experiments) in 4% paraformaldehyde (PFA) in 1✕ PBS at 4 °C before transferring to 30% sucrose in PBS at 4 °C overnight. Tissues were then washed in PBS and frozen in OCT using an ethanol/dry-ice slurry for sectioning. 10 µm thick sections were washed with PBS before incubation in 5% FBS and 0.3% Triton X-100 in PBS for 1 hour. All primary and secondary antibodies were diluted in 1% FBS and 0.3% Triton X-100 in PBS. Sections were incubated with Rabbit anti-RFP primary antibody (Rockland, 600-401-379, 1:200 dilution), Rat Anti-Mouse CD31 (BD Pharmingen™, 557355, 1:100 dilution), anti-MAP2 Antibody (Biolegend, 822501, 1:500 dilution) overnight at 4 °C before washing 3 times with PBS and staining with Goat anti-Rat IgG Alexa Fluor™ 488 (Invitrogen, A11006, 1:500 dilution), Goat anti-Rabbit IgG Alexa Fluor™ Plus 555 (Invitrogen, A32732, 1:500 dilution), and Goat anti-Chicken IgY Antibody, Alexa Fluor™ Plus 647 (Invitrogen, A32933, 1:500 dilution) secondary antibodies for 1 hour at room temperature. Sections were washed three times with PBS and stained with DAPI diluted in PBS (Thermo Scientific, 62248, 1:10,000 dilution) for 5 minutes before being washed and mounted (Molecular Probes™ ProLong™ Diamond Antifade Mountant, P36965). Slides were dried overnight before being sealed with ProLong™ Coverslip Sealant (Invitrogen, P56128).

### Quantitative PCR (qPCR) and Reverse transcription-quantitative PCR (RT-qPCR)

For RT-qPCR experiments using C57BL/6J mice, tissues were collected in RNALater (Invitrogen, AM7012) and stored in 4 °C until use. RNA was extracted using an RNeasy Mini Kit and following the manufacturer’s protocol. For RT-qPCR experiments using B-hTfR1 mice, tissues were perfusion fixed and stored in OCT as described in the previous section, and RNA was extracted as previously reported^60^. Briefly, tissues were homogenized in 200 μl of lysis buffer (10 mM Tris-HCl [pH 8.0], 0.1 mM EDTA, 0.2% SDS) and incubated at 80°C in a digital dry bath for 10 min. 200 μl of 1 mg/ml proteinase K buffer (1 mg/ml proteinase K in 10% SDS, 50 mM EDTA and 10 mM Tris-HCl (pH 7.4)) was added and samples were incubated for 15 min in a digital dry bath at 56°C. 1 ml of TRIzol reagent (Invitrogen, 15596018) was added, and samples were vortexed for 30 seconds. 300 μl of chloroform was added, and samples were again vortexed for 30 seconds before centrifugation at 13,500 RCF for 15 min at 4 °C. The clear upper aqueous phase was mixed with 1 volume of 70% ethanol before application to an RNeasy spin column, after which the standard RNeasy Mini Kit protocol was followed. All cDNA was synthesized from extracted RNA using the ProtoScript® II First Strand cDNA Synthesis Kit (New England Biolabs, E6560), as per the manufacturer’s instructions.

DNA was extracted from C57BL/6J mouse tissues using Qiagen DNeasy Blood & Tissue Kit according to the manufacturer’s protocol. For DNA extraction from fixed B-hTfR1 mouse tissues, samples were additionally incubated at 90°C for 1 hour following proteinase K digestion at 56°C. qPCR of extracted DNA and synthesized cDNA was performed using the iTaq Universal SYBR Green Supermix (Bio-Rad, 1725122) for 40 cycles with primers against either the transgene or GAPDH. Results were averaged from triplicates and were normalized to GAPDH expression using the ΔΔCq method.

### Imaging

TEM images were acquired using an FEI Tecnai Spirit G2 BioTWIN Transmission Electron Microscope. Confocal imaging was performed at the Nikon Imaging Center using the Krieger BioPipeline high-content imaging system (Nikon Eclipse Ti2-E platform with spinning disk confocal module and NIS-Elements software). Whole-slide imaging was performed at the UC San Diego School of Medicine Microscopy Core using an Olympus VS200 slide scanner (widefield automated whole-slide imaging system). Images of PBMCs were obtained using a Leica DMi8 inverted microscope equipped with an Andor Zyla sCMOS camera.

### Statistics and Reproducibility

Statistical analyses were performed using GraphPad Prism (version 10.6.0). Unless otherwise stated, data are shown as mean ± standard error of the mean (SEM). Comparisons among more than two groups were analyzed using one-way ANOVA followed by Tukey’s multiple comparisons test. Experiments involving two independent variables were analyzed using two-way ANOVA with Dunnett’s multiple comparisons test. The statistical test used for each experiment is specified in the corresponding figure legend. Significance is defined as p < 0.05.

## SUPPORTING INFORMATION

Additional details for characterization of MATCH-AAV vectors and performance of brain- and T cell-targeted MATCH-AAV-DJ vectors **(Supp. Fig. 1)**. Characterization of PBMC transduction by T cell-targeted MATCH-AAV9 vectors **(Supp. Fig. 2)**. Additional data for in-vivo administration of mTfR1-targeted MATCH-AAV9 vectors **(Supp. Fig. 3)**. Confocal microscopy images showing brain transgene expression from mice transduced with mTfR1-targeted MATCH-AAV9 vectors **(Supp. Fig. 4)**. Additional in vitro screening data and sagittal brain sections from hTfR1-targeting experiments **(Supp. Fig. 5).** Confocal microscopy images showing transgene expression in the brainstem and cerebral cortex from mice transduced with hTfR1-targeted MATCH-AAV9 vectors **(Supp. Fig. 6)**. Confocal microscopy images showing transgene expression in the midbrain and cerebellum from mice transduced with hTfR1-targeted MATCH-AAV9 vectors **(Supp. Fig. 7)**.

**Supplementary Figure 1.**
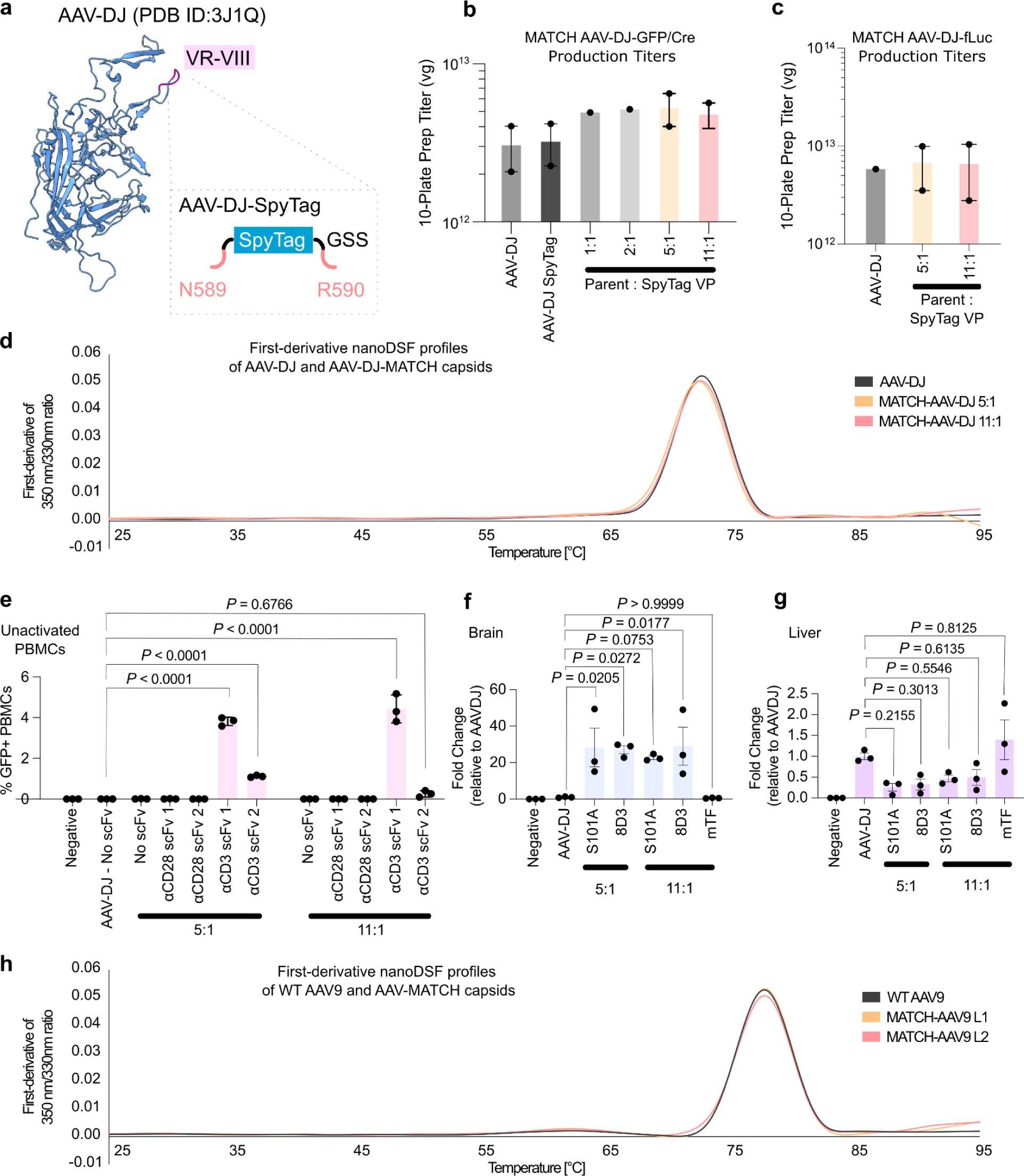
MATCH capsid characterization and MATCH-AAV-DJ transduction of resting PBMCs and mouse tissues. **(a)** Structure of AAV-DJ highlighting variable region VIII used for SpyTag insertion with GSS linker. Production titers of vectors containing either **(b)** GFP/Cre or **(c)** fLuc transgenes, composed of various AAV-DJ-VP:AAV-DJ-SpyTag-VP ratios. **(d)** nanoDSF profiles of profiles of AAV-DJ, MATCH-AAV-DJ 5:1, and MATCH-AAV-DJ 11:1 capsids containing a fLuc transgene. **(e)** Transduction of resting PBMCs using mosaic SpyTag-AAV-DJ variants conjugated to αCD3 or αCD28 scFvs at an MOI of 1 ✕ 10^4^ vg/cell. **(f)** Brain and **(g)** liver luciferase expression three weeks after retro-orbital delivery of 1 ✕ 10^12^ vg MATCH-AAV-DJ conjugated to S101A or 8D3 αmTfR1-scFv-SpyCatcher. Data represent mean ± SEM. One-way ANOVA with Tukey’s multiple comparisons test **(e–g)**. **(h)** nanoDSF profiles of WT AAV9, MATCH-AAV9 L1, and MATCH-AAV9 L2 capsids containing an mCherry transgene.

**Supplementary Figure 2.**
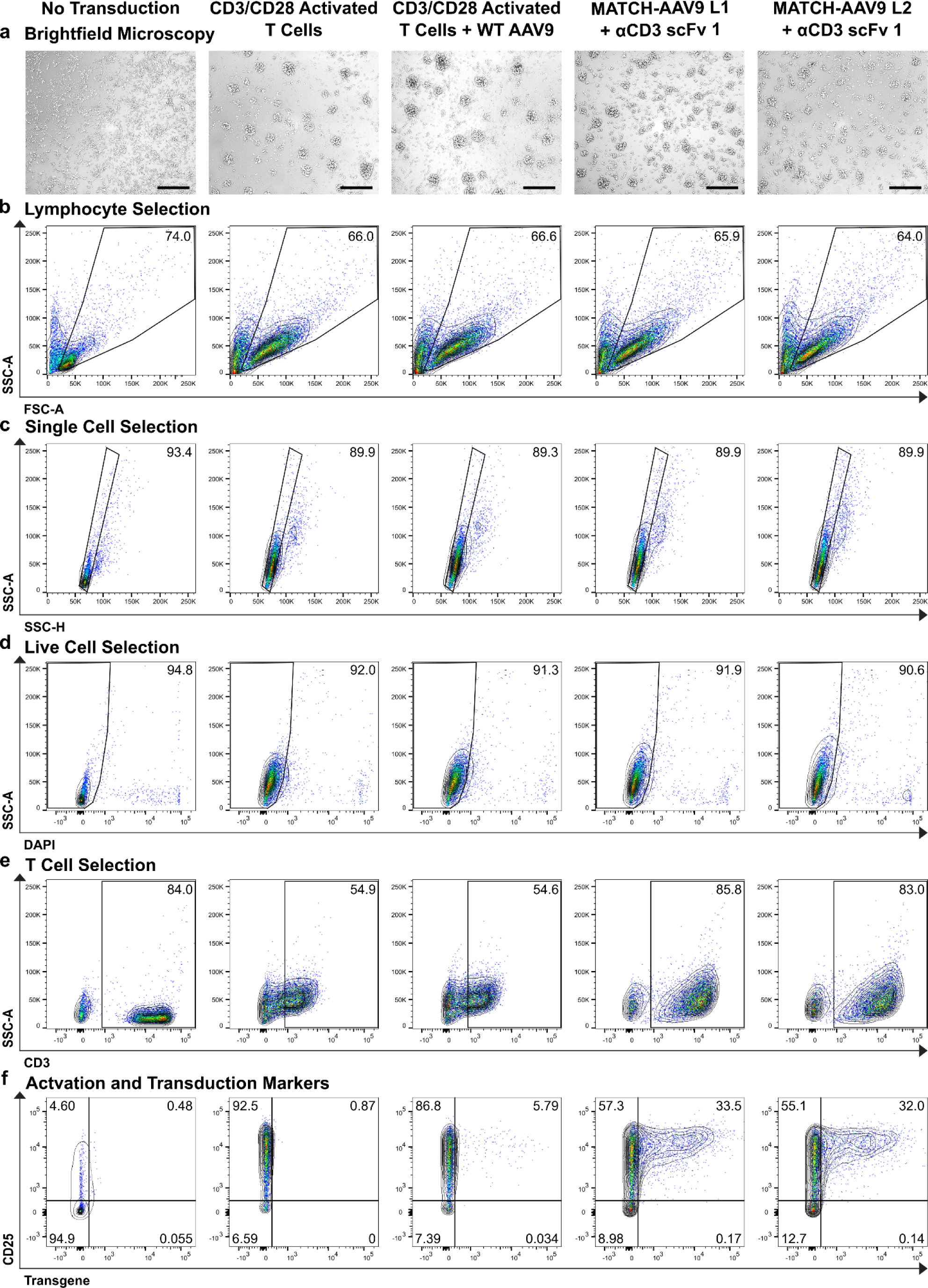
CD3-targeted MATCH-AAVs activate and transduce T cells. **(a)** Microscopy of PBMC cultures which were untransduced or transduced with MATCH-AAV9 conjugated to αCD3-scFv 1 at an MOI of 1 ✕ 10^5^ vg/cell. Scale Bar 250 μm. **(b)** Flow cytometry gating expanded to include activated lymphocyte populations. **(c)** Gating for single cell and **(d)** live cell selection. **(e)** Selection of CD3⁺ T cells. **(f)** Expression of CD25 activation marker and mCherry transgene in transduced T cells.

**Supplementary Figure 3.**
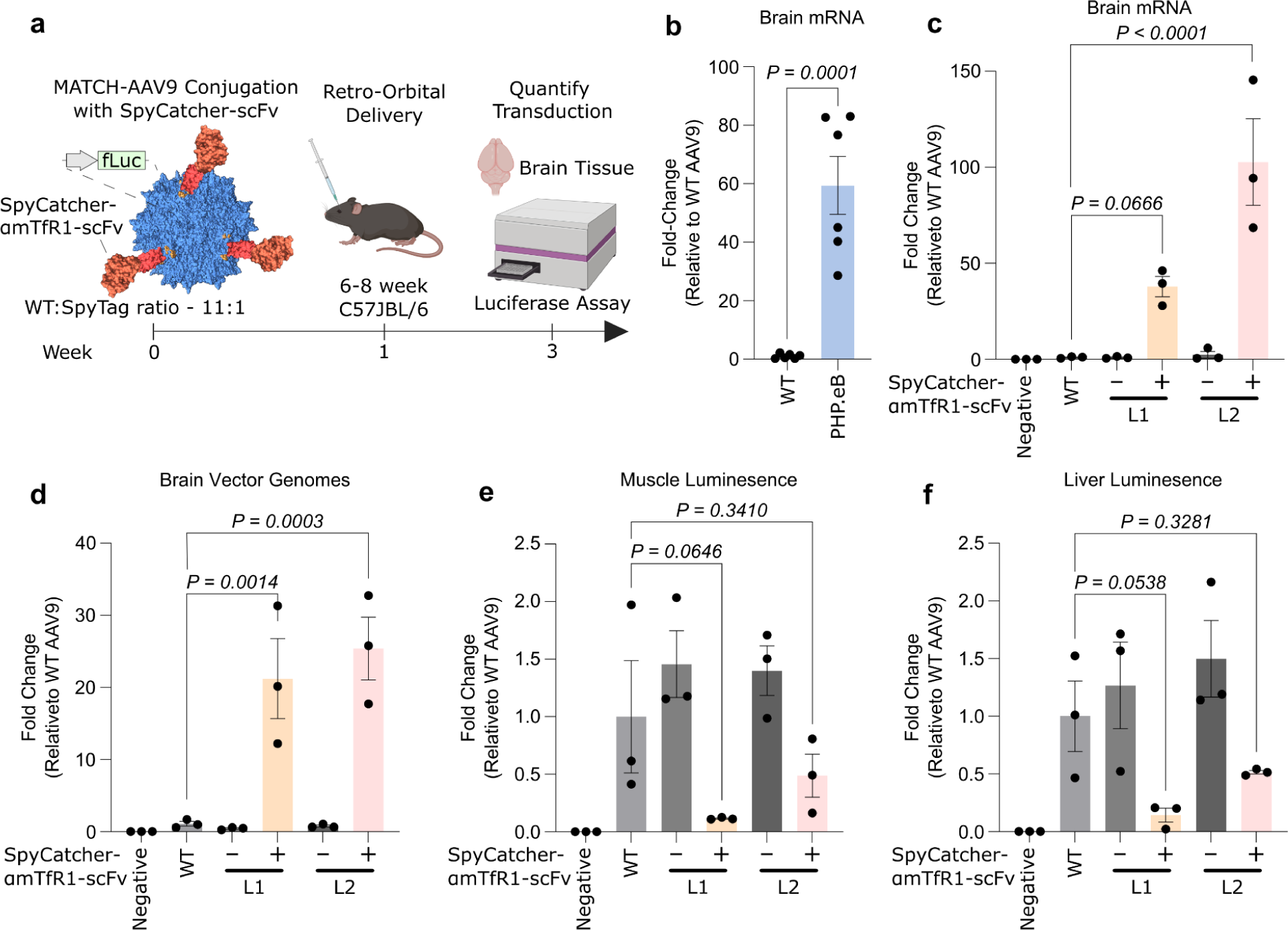
Transduction profile of MATCH-AAV9 variants in-vivo and in HEK293T cells relative to AAV9-WT. **(a)** Workflow for in vivo CNS transduction using MATCH-AAV9 conjugated to αmTfR1-scFv-SpyCatcher. Prevalence of **(b)** PHP.eB and **(c)** MATCH-AAV9 mRNA transcripts in the murine brain relative to AAV9 three weeks after retro-orbital delivery of 1 ✕ 10^12^ vg as determined by RT-qPCR**. (d)** Prevalence of MATCH-AAV9 vector genomes in the murine brain relative to AAV9 three weeks after retro-orbital delivery of 1 ✕ 10^12^ vg as determined by qPCR. Fold improvement in **(e)** muscle and **(f)** liver luciferase expression over WT AAV9, three weeks after retro-orbital injection of 1 ✕ 10^12^ vg. Data represent mean ± SEM. One-way ANOVA with Tukey’s multiple comparisons test **(b–f)**.

**Supplementary Figure 4.**
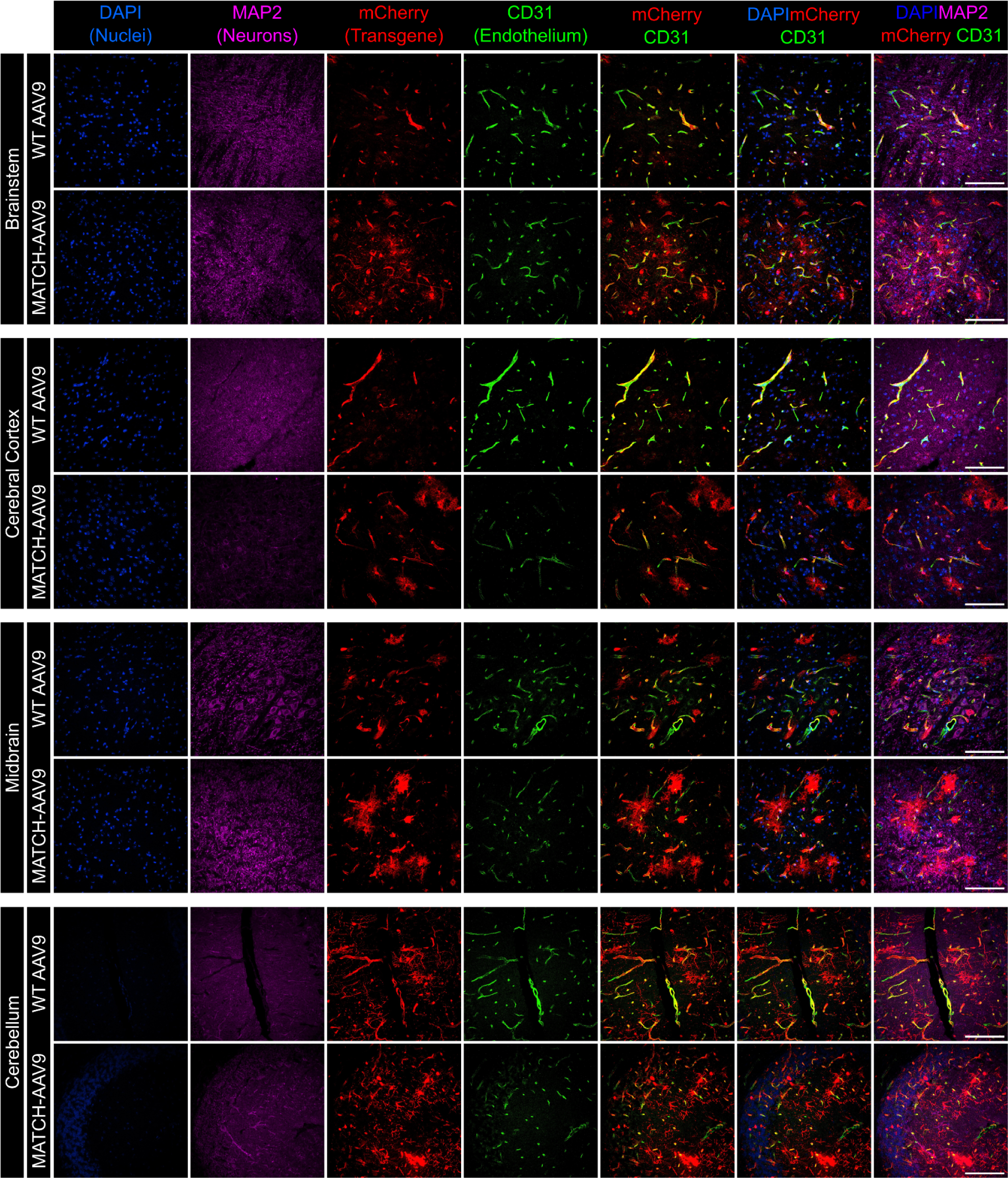
MATCH-AAV9 L2 crosses the BBB and enables CNS-wide transduction in-vivo. Images of C57BL6/J mouse brain sections three weeks after retro-orbital delivery of 5 ✕ 10^12^ vg/mouse WT AAV9 or MATCH-AAV9 L2 conjugated to scFv-8D3 with nuclei (DAPI, blue), MAP2 (neuronal cytoplasm, magenta), mCherry transgene (red), and endothelium (CD31, green) labeled via immunohistochemistry. Scale Bar 100 μm.

**Supplementary Figure 5.**
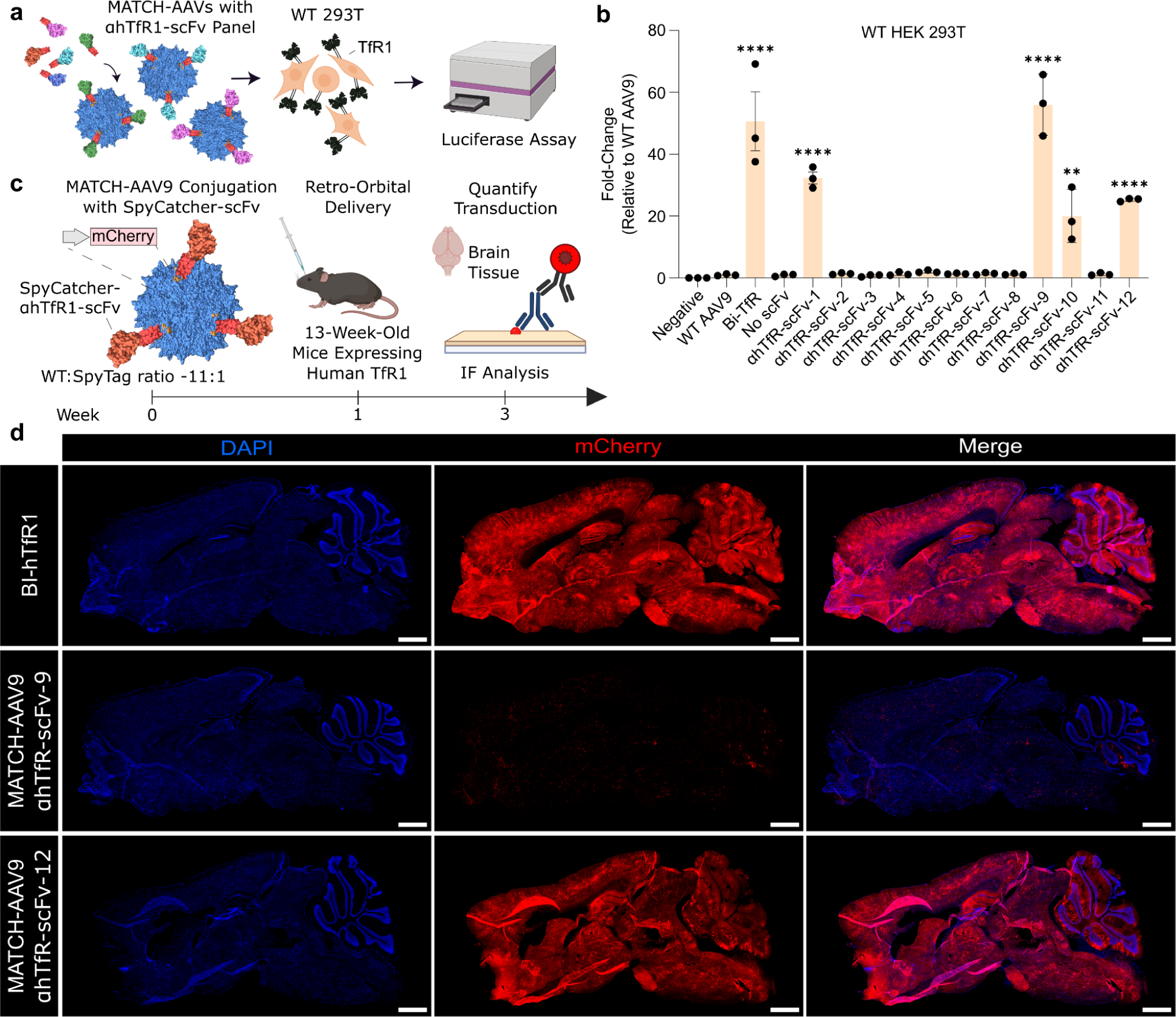
MATCH-AAV9 L2 enables efficient hTfR1-targeted transduction of WT HEK293Ts and the brain. **(a)** Workflow for transduction of WT HEK293T cells with αhTfR1-scFv-SpyCatcher-decorated MATCH-AAV9 L2. **(b)** Evaluation of transferrin-mediated transduction by αhTfR1-scFv-SpyCatcher-decorated MATCH-AAV9 L2 in WT HEK293T cells at an MOI of 1 ✕ 10^4^ vg/cell. Data represent mean ± SEM. **P < 0.01, ****P < 0.0001 (one-way ANOVA with Tukey’s multiple comparisons test). **c)** Workflow for in vivo CNS transduction using MATCH-AAV9-L2 conjugated to αhTfR1-scFv-9 or αhTfR1-scFv-12. **(d)** Sagittal brain sections immunostained for nuclei (DAPI) and AAV transgene (mCherry) from mice three weeks after retro-orbital delivery of 1 ✕ 10^12^ vg/mouse BI-hTfR1 or MATCH-AAV9 conjugated to αhTfR1-scFv-12. Scale Bar 1 mm.

**Supplementary Figure 6.**
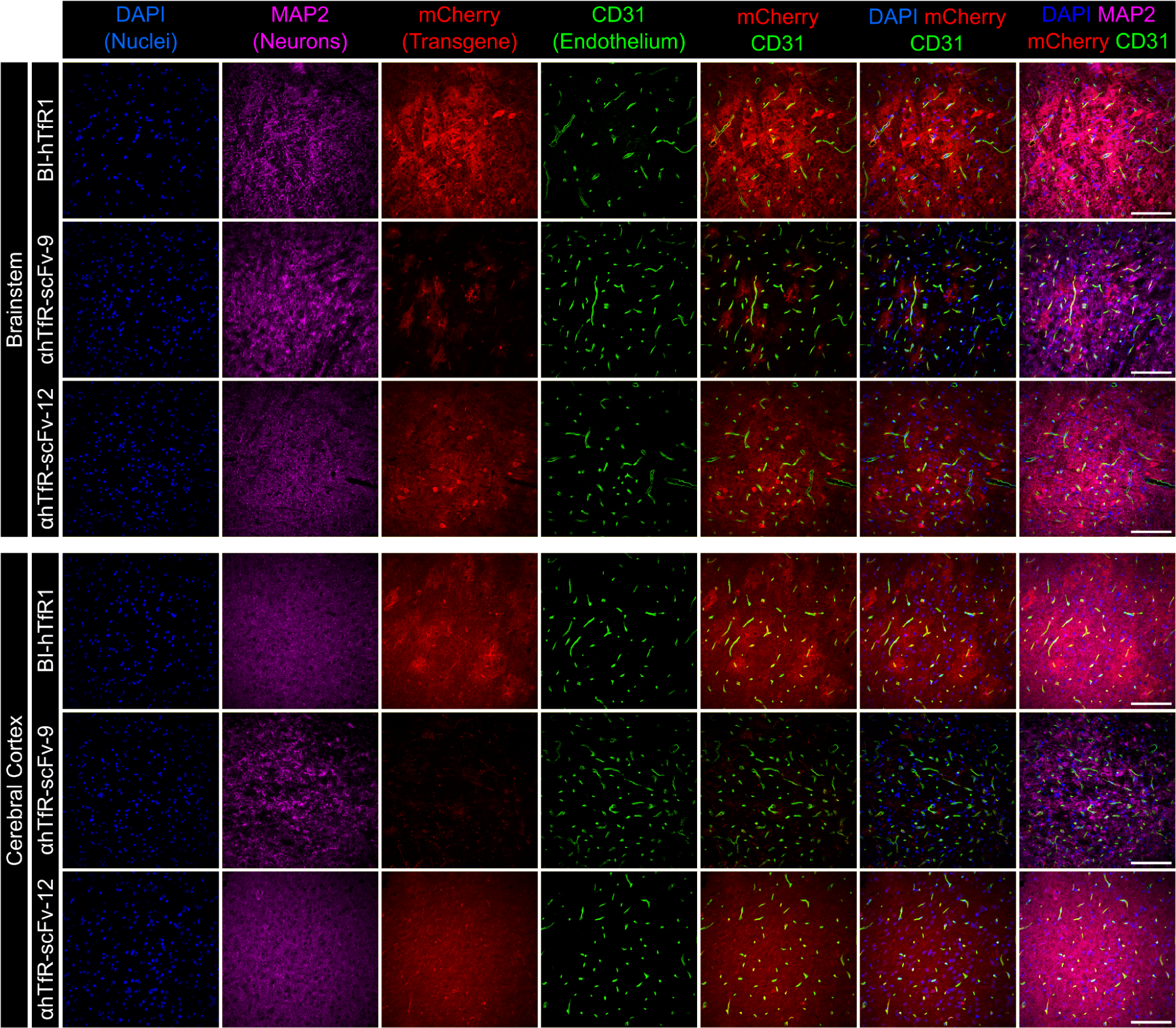
MATCH-AAV9 L2 crosses the BBB in the brainstem and cerebral cortex of humanized mice in-vivo. Images of B-hTFR1 mouse brain sections from the brainstem and cerebral cortex, transduced with either 1 ✕ 10^12^ vg/mouse BI-hTfR1 or MATCH-AAV9 L2 conjugated to αhTfR1-scFv-9 or αhTfR1-scFv-12 with nuclei (DAPI, blue), MAP2 (neuronal cytoplasm, magenta), mCherry transgene (red), and endothelium (CD31, green) labeled via immunohistochemistry. Scale Bar 100 μm.

**Supplementary Figure 7.**
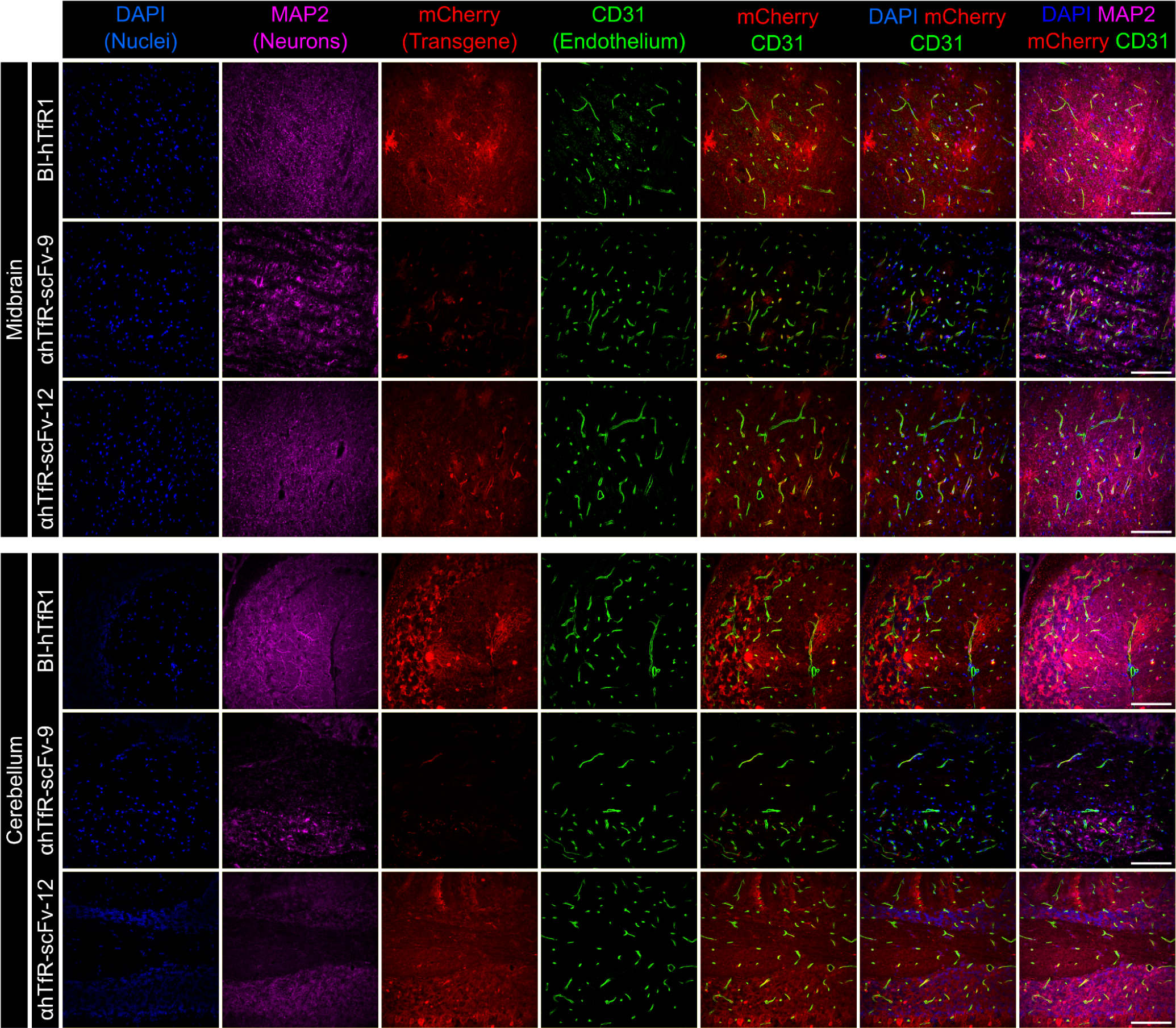
MATCH-AAV9 L2 crosses the BBB in the midbrain and cerebellum of humanized mice in-vivo. Images of B-hTFR1 mouse brain sections from the midbrain and cerebellum, transduced with either 1 ✕ 10^12^ vg/mouse BI-hTfR1 or MATCH-AAV9 L2 conjugated to αhTfR1-scFv-9 or αhTfR1-scFv-12 with nuclei (DAPI, blue), MAP2 (neuronal cytoplasm, magenta), mCherry transgene (red), and endothelium (CD31, green) labeled via immunohistochemistry. Scale Bar 100 μm.

